# Dopaminergic drugs decrease loss aversion in Parkinson’s disease with but not without depression

**DOI:** 10.1101/069047

**Authors:** Monique H.M. Timmer, Guillaume Sescousse, Rianne A.J. Esselink, Payam Piray, Roshan Cools

## Abstract

Depression, a common non-motor symptom of Parkinson’s disease (PD), is accompanied by impaired decision making and an enhanced response to aversive outcomes. Current strategies to treat depression in PD include dopaminergic medication. However, their use can be accompanied by detrimental side effects, such as enhanced risky choice. The mechanisms underlying dopamine-induced increases in risky choice are unclear. In the current study we adopt a clinical-neuroeconomic approach to investigate the effects of dopaminergic medication on loss aversion during risky choice in depressed and non-depressed PD. Twenty-three healthy controls, 21 depressed and 22 non-depressed PD patients were assessed using a well-established gambling task measuring loss aversion during risky choice. Patients were tested on two occasions, after taking their normal dopaminergic medication (ON) and after withdrawal of their medication (OFF). Dopaminergic medication decreased loss aversion to a greater extent in depressed than non-depressed PD patients. Moreover, we show that the degree to which dopaminergic medication decreases loss aversion correlated with current depression severity and with drug effects on depression scores. These findings demonstrate that dopamine-induced changes in loss aversion depend on the presence of depressive symptoms in PD.

**Significance statement:** Dopaminergic medication that is used to treat motor and non-motor symptoms in patients with Parkinson’s disease is known to contribute to risky decision-making. The underlying mechanisms are unclear. The present study demonstrates that dopaminergic medication in Parkinson’s disease decreases loss aversion during risky choice, but only in depressed and not in non-depressed patients with Parkinson’s disease. These results advance our understanding of the mechanisms underlying dopamine-induced risky choice, while also identifying depression as an important factor that confers vulnerability to such dopamine-induced risky choice.

*Conflict of Interest:* The authors declare no competing financial interests.

## Introduction

Depression is a common non-motor symptom of Parkinson’s disease (PD) which greatly affects quality of life (Schrag, 2006). Similar to the motor symptoms, depression in PD can be treated with dopaminergic medication (Barone et al., 2010; Stacy et al., 2010; Seppi et al., 2011). However, their use is limited by potential side effects, such as enhanced risk-taking behavior, in their most severe form qualifying as impulse control disorder (ICD) (Weintraub et al., 2010; Voon et al., 2011b). The mechanisms underlying dopamine-induced increases in risky choice have remained unclear.

One mechanism by which dopaminergic medication can increase risky choice is by attenuating loss aversion. Loss aversion reflects the relative weighting of gains and losses during risky choice and is one of the core concepts of Prospect Theory, a well-known economic theory of decision-making under risk (Kahneman and Tversky, 1979, 1984). In the domain of learning, dopamine manipulation studies in healthy controls and PD patients have revealed that the balance between learning from reward and punishment critically depends on striatal dopamine (Frank et al., 2004; Cools et al., 2006; van der Schaaf et al., 2014). Increases in dopamine enhance reward-based learning/choice while impairing punishment-based learning/choice and decreases in dopamine enhance punishment-based learning/choice while impairing reward-based learning/choice. One obvious next question is whether dopaminergic medication alters risky choice in an analogous manner, by increasing the weighting of prospective rewards (gains) relative to punishments (losses). This relative weighting corresponds exactly to loss aversion in the context of Prospect Theory.

Depression has been associated with reduced reward and enhanced punishment sensitivity across various domains including decision making (Eshel and Roiser, 2010). For instance, depressed individuals (without PD) have been shown to exhibit reduced reward-based reversal learning and attenuation of associated BOLD signal in the ventral striatum (Robinson et al., 2011). Depressed patients also exhibit enhanced loss-minimization and attenuated gain-maximization as well as enhanced loss aversion during risky choice (Gradin et al., 2011; Maddox et al., 2012; Chandrasekhar Pammi et al., 2015). This cognitive profile, together with clinical observations that depressive symptoms in PD occur more often during OFF periods (Maricle et al., 1998) and can (to some degree) be alleviated by dopaminergic medication (Barone et al., 2010), concurs with evidence indicating that depression in PD is associated with dopamine deficiency in the ventral striatum (Weintraub et al., 2005; Vriend et al., 2013; Vriend et al., 2014).

Based on current available evidence, we put forward two opposite hypotheses about the effects of dopaminergic medication on loss aversion in depressed PD patients. According to one account of drug-induced cognitive deficits, the “dopamine overdose hypothesis”, dopaminergic doses necessary to remedy the severely dopamine depleted dorsal striatum and associated cognitive and motor functions, might detrimentally overdose relatively intact ventral striatal dopamine levels (Gotham et al., 1988; Swainson et al., 2000; Cools et al., 2001). One implication of this hypothesis is that drug-induced increases in risky choice are restricted to patients with intact ventral striatal dopamine levels, while not extending to patients with already depleted ventral striatal dopamine levels, such as those with co-morbid depression. The alternative hypothesis stems from clinical observations showing that specifically depressed PD patients are more likely to experience ICDs (Isaias et al., 2008; Joutsa et al., 2012), suggesting that dopamine-induced increases in risky choice are restricted to depressed patients.

To disentangle these hypotheses, we compared PD patients, with and without depression on a gambling task measuring loss aversion on two occasions: once ON and once OFF dopaminergic medication.

## Materials & Methods

### Participants and experimental design

We recruited 23 non-depressed PD patients, 24 depressed PD patients and 25 healthy controls. Data from 1 non-depressed patient, 3 depressed patients and 2 healthy controls were discarded from analyses for several reasons (see exclusion). Patients were recruited from the Parkinson Centre at the Radboud university medical centre, the Netherlands. Healthy controls were recruited via advertisement, or were partners or acquaintances of patients. Healthy controls and patients were matched for gender, age and IQ measured with the NART (Dutch version of the National Adult Reading Test (Schmand et al., 1991)). Furthermore, patient groups were matched in terms of disease severity (measured with the Unified Parkinson’s Disease Rating Scale (UPDRS part III) (Goetz and Stebbins, 2004)) and used similar amounts of dopaminergic medication (LED (Levodopa Equivalent Dose (Esselink et al., 2004))(Table 1). Written informed consent according to the Declaration of Helsinki was obtained from all participants. The study was part of a larger project investigating the neurobiological mechanisms of depression in PD and was approved by the local ethics committee (CMO region Arnhem -Nijmegen, the Netherlands, nr. 2012/43).

**Table 1.** Group characteristics

|  | Depressed PD | Non-depressed PD | Healthy controls |
| --- | --- | --- | --- |
|  | n = 21 | n = 22 | n = 23 |
| Gender, men | 13 | 13 | 14 |
| Age, years | 58.5 (5.8) | 61.0 (7.6) | 60.9 (5.9) |
| NART-IQ | 96.2 (11.6) | 97.0 (15.5) | 100.7 (13.7) |
| MMSE | 28.5 (1.4) | 28.6 (1.3) | 28.8 (1.2) |
| Hoehn & Yahr | 1.6 (0.4) | 1.8 (0.5) | - |
| UPDRS - III (OFF) | 22.7 (9.6) | 22.2 (6.5) | - |
| Disease duration, years | 5.1 (3.5) | 4.5 (2.2) | - |
| LED mg/day | 551 (248) | 627 (275) | - |
| LED agonists mg/day | 71 (122) | 103 (129) | - |
| BDI (OFF) | 9.9 (6.1) | 4.0 (2.3) | 3.1 (2.1) |
| Current ICD | 4 | 1 | - |
| First session ON | 11 | 9 | - |
| Days between sessions | 23 (27) | 21 (20) | - |
| LED = Levodopa Equivalent Dose (Esselink et al., 2004) |  |  |  |

Values represent numbers or mean (standard deviation)

All patients were diagnosed with idiopathic PD according to the UK Brain Bank criteria (Gibb and Lees, 1988) by a neurologist specialized in movement disorders (Prof. B.R. Bloem, Dr. R.A. Esselink, Dr. B. Post) and were treated with dopaminergic medication. In the non-depressed patient group 11 patients were treated with levodopa, 2 with dopamine receptor agonists and 9 with both. In the depressed patient group, 14 patients were treated with levodopa, 2 with dopamine receptor agonists and 5 with both. Moreover, 7 depressed patients received antidepressants (paroxetine n=3, escitalopram n=1, venlafaxine n=1 and nortriptyline n=2). Patients were on stable medication regimes during the course of the study, except for one patient who used duloxetine – a serotonin/noradrenalin reuptake inhibitor prescribed to treat pain - for 4 weeks between the two testing days (in this case testing days were separated by 17 weeks). The drug was discontinued 4 weeks before the second testing day.

Patients were included in the depressed group if they met the DSM-IV criteria for a major (n=7) or minor depressive episode (n=12), dysthymic disorder (n=1) or adjustment disorder with depressed mood (n=1) within five years before PD diagnosis up until now. This five-year cut-off was chosen because the incidence of depression is significantly higher within the five years before PD diagnosis and therefore likely related to PD pathology (Shiba et al., 2000). Thus, PD patients were selected based on a PD-related depression (history) rather than current depressive symptoms. Seven patients were identified as having current depression. Psychiatric diagnosis was based on structured psychiatric interviews administered during an intake session (MINI-plus (Sheehan et al., 1998)). General exclusion criteria were clinical dementia (Mini Mental State Examination < 24, (Folstein et al., 1975)), psychiatric disorders other than depression (bipolar disorder, schizophrenia, ADHD and drug or alcohol abuse), neurological co-morbidity and hallucinations. Healthy controls were also excluded if they had a history of mood or anxiety disorder, obsessive-compulsive disorder or used any psychotropic medication.

Patients were assessed on two occasions, once after taking their normal dopaminergic medication (ON) and once after abstaining from their dopaminergic medication for at least 18 hours (24 hours for slow release dopamine receptor agonists) (OFF). Patients who used antidepressants were asked to take these antidepressants on both testing days enabling us to assess specifically dopaminergic drug effects on gambling behavior. The order of ON and OFF sessions was counterbalanced in each patient group (Table 1). Healthy controls were only tested once. During testing sessions we administered the gambling task described below. Furthermore, on each testing day, participants completed the Beck Depression Inventory (BDI (Beck et al., 1961)) to assess current depressive symptoms. Participants were instructed to answer BDI questions, not according to how they felt over the past week, but according to how they felt over the past 24 hours, enabling us to assess dopaminergic drug (withdrawal) effects on depression scores. Patients also completed the QUIP rating scale (Weintraub et al., 2012) developed to assess ICD symptoms in PD and were clinically assessed on motor symptom severity (UPDRS part III (Goetz and Stebbins, 2004)).

Participants were paid a fixed amount per testing day for participation (healthy controls; 30 Euros, patients; 40 Euros) and received an additional amount of money based on task performance (between 2-11 Euros per session).

### Task

Participants played a well-validated gambling task designed to measure loss aversion (Figure 1) (Tom et al., 2007). During this task, participants were presented with 169 mixed gambles (split into 3 runs) on a computer screen. Each gamble offered a 50/50 percent chance of either gaining or losing varying amounts of money. Potential gains ranged from +€6 to +€30 (increments of €2), potential losses ranged from -€3 to -€15 (increments of €1). This asymmetric gain-loss range was chosen in order to maximize statistical power, based on the assumption that on average people are twice as sensitive to losses as they are to gains (Tom et al., 2007). Each of the possible gain-loss pairs (13x13=169) was presented once in randomized order. Participants were asked to either accept (play) or reject the gamble by pressing one of two buttons. In order to make participants feel that they were gambling with *their own money*, and thus avoid “house money effects” (Thaler and Johnson, 1990), endowments at the beginning of this gambling task were earnings from a behavioral experiment immediately preceding the present experiment on the same day. Gambles were not resolved during the experiment to exclude behavioral adjustments on a trial-by-trial basis. However, participants were told to take each gamble seriously, because at the end of the experiment, 3 gambles would be randomly selected and played for real money.

**Figure 1.**
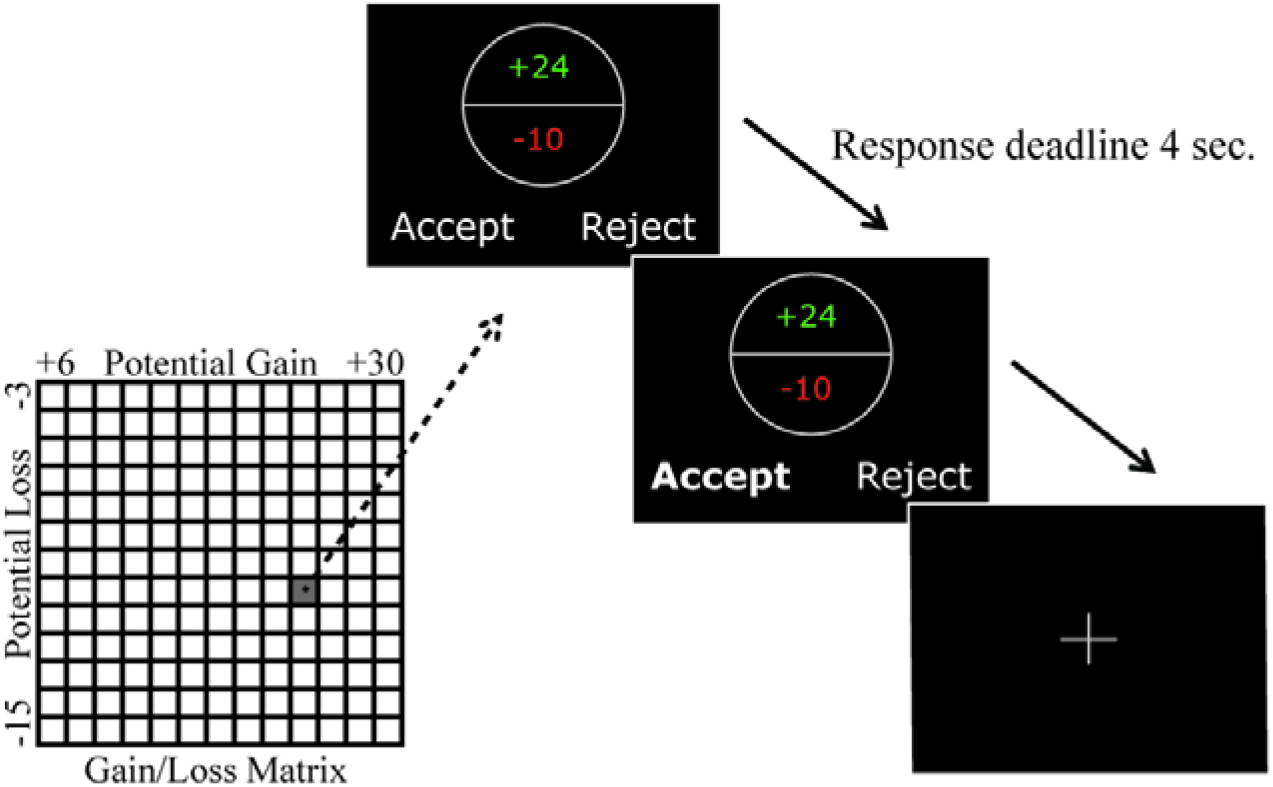
Task overview. Participants played a gambling task designed to measure loss aversion. During this task participants were presented with 169 mixed gambles, each offering 50/50 percent change of either gaining or losing varying amounts of money. Gains ranged from +€6 to +€30 (increments of €2), losses ranged from -€3 to - €15 (increments of €1) (see gain/loss matrix). Each possible gain/loss pair was presented once in randomized order. Participants were asked to either accept (play) or reject the gamble within a maximum time of 4 sec.

## Analysis

### Model

We used a model-based approach to analyze participant’s choice behavior. This procedure involved fitting a theoretical model of decision making to the behavioral data in order to quantify specific aspects of choice behavior. One of the most popular accounts of decision making under risk is Prospect Theory (Kahneman and Tversky, 1979). We sought to understand the effects of dopaminergic medication and depression in PD in light of this theory by assessing effects of medication and PD-related depression diagnosis on parameters obtained from a model based on Prospect Theory. Within that framework, the subjective utility of each gamble (SUG) can be approximated by the following equation:

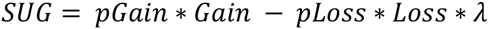

Where p_Gain_ is the gain probability, p_Loss_ the loss probability, *Gain* the gain value of the gamble and *Loss* the (absolute) loss value of the gamble. The relative weighting of gains and losses is reflected in the loss aversion parameter λ. If λ>1, then losses are overvalued relative to gains: a person is loss averse. If λ<1, then gains are overvalued relative to losses: a person is loss seeking. And if λ=1, gains and losses are valued equally: a person is gain-loss-neutral.

A softmax function was used to estimate the probability of gamble acceptance based on the subjective value of the gamble:

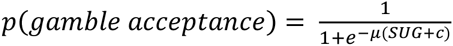

Using this procedure we obtained two other parameters: the inverse temperature parameter (μ) and a constant parameter (c). The inverse temperature parameter reflects consistency of choice behavior. If μ is zero, choices are random, whereas if μ is highly positive or negative, there is consistency in choice behavior, with a positive μ representing higher gamble acceptance with higher gain and lower loss value (and vice versa for negative μ). We anticipated μ to be positive, consistent with a utility maximization strategy, where participants accept more gambles when gain values increase and loss values decrease. The constant parameter (c) reflects a response bias toward or away from gambling irrespective of the value of the gambles. If c>0, there is a tendency to accept gambles regardless of their subjective utility. If c<0, there is a tendency to reject gambles regardless of their subjective utility.

The model that we fitted to the data assumes a linear valuation of gains and losses, in contrast to the curvilinear value function of Prospect Theory. This is a common and reasonable simplifying assumption given the relatively narrow range of gains and losses used in this protocol. We also assumed no subjective transformation of probabilities as described in Prospect Theory and thus assumed equal weights for the 0.5 probability of gains and losses (Tom et al., 2007; De Martino et al., 2010).

### Exclusion

We assessed whether participants’ choices were influenced by gain and loss values in an expected manner, i.e. whether participants were utility maximizers (accepting more gambles with increasing gain values and accepting fewer gambles with increasing loss values). Inspection of the individual responses revealed that two participants (one depressed and one non-depressed PD patient) did not meet this a priori assumption, suggesting a lack of understanding of task instructions. In both cases this was during the first testing day. In one case the response graph revealed that the participant accepted more gambles when gain values decreased and loss values increased, thereby unintentionally trying to minimize earnings. During debriefing this participant realized that he had made a mistake. The responses of the other participant were suggestive of random choice behaviour. In both cases, these observations were confirmed by negative temperature parameters (μ) obtained from the model. These two patients were excluded from further analyses. Moreover, two healthy controls were excluded from further analyses because of a lifetime history of depression, while two depressed PD patients were excluded because they failed to finish the study leading to incomplete datasets. The final analysis included 23 healthy controls, 22 non-depressed PD patients and 21 depressed PD patients.

### Model fitting and comparison

We used a hierarchical Bayesian fitting procedure to fit the model to participants’ choices as described by Huys et al. (Huys et al., 2011; Huys et al., 2012). This method estimates the mean and the variance of model parameters across all subjects and sessions. These prior parameters then serve to define a normal priori distribution for finding individual values of parameters for each subject and session (i.e. posterior parameters). We hypothesized the *a priori* distribution of the relevant parameter (i.e. the loss aversion parameter λ) to be different for patients and healthy controls. Therefore we first fitted the model to patient data only. Note that any differences in posterior parameters between patient groups and medication sessions cannot be attributed to parameter regularization employed during fitting, because individual parameters from both patient groups (depressed and non-depressed) and both sessions (ON and OFF medication) were obtained using the same a priori distribution (Huys et al., 2012). In a subsequent step, to compare PD patients with healthy controls, we fitted the model to healthy control and patient data together (separately for each drug session).

A Bayesian model comparison was conducted to compare the model with 3 parameters (λ, μ and c) with a slightly simpler model, where we forced c to be zero, thereby reducing the number of free parameters. This model assumed that subjects do not exhibit a response bias toward or away from gambling irrespective of the value of the gambles. A Bayesian model comparison assessed which model best captured participants’ choices by computing model evidence by balancing model fits and model complexity (Kass and Raftery, 1995; Piray et al., 2014)(MacKay et al., 2013). A procedure was employed that penalizes complexity by marginalizing over both group and individual parameters using Laplace approximation and Bayesian information criterion, respectively. The negative log-mode evidence (NLME) was computed as:

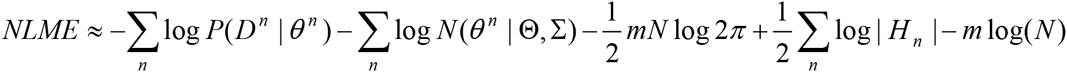

where *D^n^* is the set of choice data for the *n*th participant, *θ^n^* is the fitted individual parameter for *n*th participant, ^^ and ^Σ^ are the mean and variance for the group distribution, respectively, ^*m*^ is the number of free parameters of the model, ^*N*^ is the number of participants and ^|*H_n_*|^ is the determinant of the Hessian matrix of the log-posterior function at ^*θ^n^*^. The first term on the right hand-side of the equation refers to how well the model predicts data. The sum of the next three terms together is the penalty due to individual parameters. The last term represents the penalty approximated for ^2*m*^ (mean and variance together) group parameters using Bayesian information criterion (Piray et al., 2014). The model with the lowest log-model evidence is the best model.

### Statistical analysis

The primary parameter of interest was the loss aversion parameter (λ). First we compared depressed PD patients with non-depressed PD patients. Subsequently we compared healthy controls with PD patients (each group and drug session separately). For normally distributed data, we used a mixed ANOVA with drug as within-subject and group as between-subject factor. For non-normally distributed data (Shapiro-Wilk, *p*<0.05) we used two-tailed Wilcoxon signed-rank tests to assess within-subject differences and Mann Whitney tests to assess between-group differences. Two-tailed Pearson correlations were used for normally distributed data and two-tailed Spearman correlations for non-normally distributed data. Furthermore, for non-normally distributed data we reported medians and their standard error. Standard errors of the median were computed using Bootstrapping (Efron et al., 1993). By resampling with replacement of the original group sample, we created 10^5^ new group samples. The standard error of the median was then defined as the standard deviation of all bootstrapped samples.

## Results

### Patient and disease characteristics

Mixed ANOVA of depression scores (BDI) from the PD patients demonstrated a significant group*drug interaction, *F*_(1,41)_=4.19, *p*=0.047. Post-hoc paired samples t-test revealed that this interaction was due to a significant drug-induced decrease in depression scores in depressed patients (*t_(20)_*=2.19, *p*=0.041) but not in non-depressed patients (*t^(21)^*=-.60, *p*=0.56). There was also a main effect of group, *F_(1,41)_*=17.26, *p*<0.001, indicating significantly higher depression scores in the depressed patient group. There was no main effect of drug (Figure 2A).

**Figure 2.**
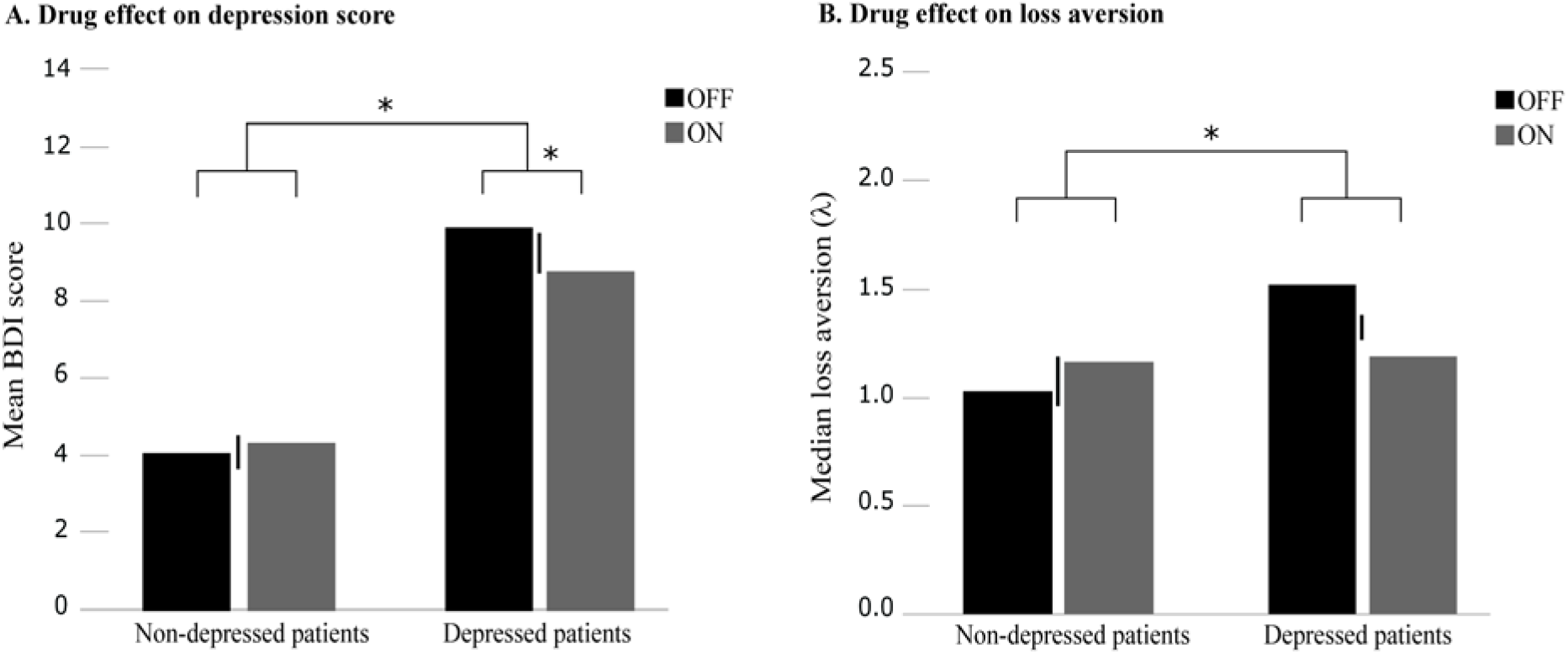
Drug effects per group and drug session. **A** Mean scores on the Beck Depression Inventory (BDI) per group (depressed and non-depressed patients) and session (OFF session in black, ON session in grey). Error bars represent standard errors of the mean difference. **B** Median loss aversion parameter (λ) per group (depressed and non-depressed patients) and session (OFF session in black, ON session in grey). Error bars represent standard errors of the median difference. \**p*<0.05

Five patients exhibited at least one ICD as assessed with the QUIP rating scale (4 depressed and 1 non-depressed patient) but the proportion of ICD was not different between the two patient groups (Chi^2^ test, *p*=0.14). None of them exhibited pathological gambling. Individual endowments at the beginning of the task varied between participants, as these were earnings from a previous experiment performed on the same day. However, there was no significant main effect of group or drug and no group*drug interaction on these earnings.

### Effects of dopaminergic drugs on loss aversion

Using Prospect Theory-based analysis, we assessed the computational mechanisms contributing to risky choice. The full model including a constant parameter (c) (reflecting a gambling response bias irrespective of the value of gambles) provided a better account of participants’ choices than did a model without this c parameter, indicated by a lower log-model evidence (in patients: 4102 compared with 4374 for the model where (c) was forced to be zero, in healthy controls: 1099 compared with 1131 for the model where (c) was forced to be zero). Therefore, reported results are based on the loss aversion parameter (λ) obtained from the full model.

The median loss aversion parameter per group and drug session can be found in Figure 2B. The loss aversion parameter (λ) was not normally distributed as indicated by Shapiro-Wilk test. Therefore we used nonparametric statistics. Our analyses revealed a significant group*drug interaction (*U*=149, *p*=0.046), which was due to greater drug-induced decreases in loss aversion in depressed patients than in non-depressed patients. If anything, medication increased loss aversion in non-depressed patients. The simple main effects of drug were not significant. There was a near-significant effect of group in the OFF state; depressed patients tended to be more loss-averse than non-depressed patients (*U*=151, *p*=0.052). During the ON state there was no group effect (*U*=215, *p*=0.70). There was no overall main effect of group (*U*=191, *p*=0.33) and no overall main effect of drug (*Z*=-0.21, *p*=0.84) (Figure 2B). There were no effects of session order.

To visualize drug and group effects on loss aversion, we plotted, for each group and drug session separately, the degree to which the ratio of rejecting to accepting gambles increased as a function of increases in potential losses (Figure 3). To control for the effects of other factors, such as general drug effects on gambling rate, we plotted the ratio of rejecting to accepting gambles as a function of relative loss differences between pairs of trials, while the effects of different gains were averaged out. A steeper slope indicates greater loss sensitivity. From this Figure 3 it is clear that dopaminergic medication had contrasting effects on loss aversion in depressed and non-depressed PD patients.

**Figure 3.**
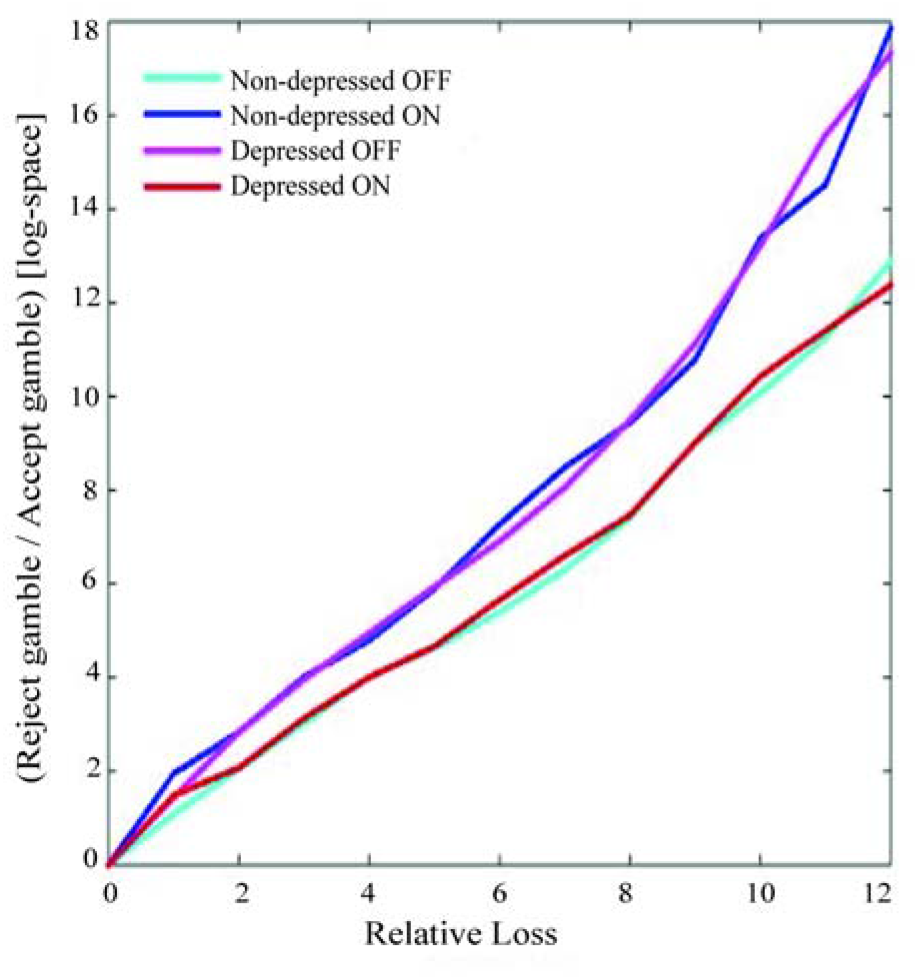
Loss sensitivity. The ratio of the number of rejected gambles divided by the number of accepted gambles in log-space (y-axis) as a function of the relative loss averaged across different gain values (x-axis) per group and per drug session. A steeper slope indicates greater loss sensitivity.

To assess whether drug effects on loss aversion were predicted by current OFF state depression severity, we performed Spearman correlations with BDI scores. Across the whole group there was a significant correlation (*rho*_(41)_=−.348, *p*=0.022). This correlation was due to greater drug-induced decreases in loss aversion in patients with higher depression scores (Figure 4A). Additionally, we investigated whether drug effects on depression scores correlated with drug effects on loss aversion. Across the whole group there was a significant correlation (*rho*_(41)_=−.384, *p*=0.011), indicating greater drug-induced decreases in loss aversion in patients with greater drug-induced decreases in depression scores. This correlation was strong in depressed patients (*rho*_(19)_=−.592, *p*=0.005), but not significant in the non-depressed patients (*rho*_(20)_=−.021, *p*=0.93) and significantly different between groups (Fisher r-z transformation, *z*=−2.01, *p*=0.044) (Figure 4B). There was no significant correlation between LED and drug effects on loss aversion (across the two groups, *rho*_(41)_=0.186, *p*=0.23).

**Figure 4.**
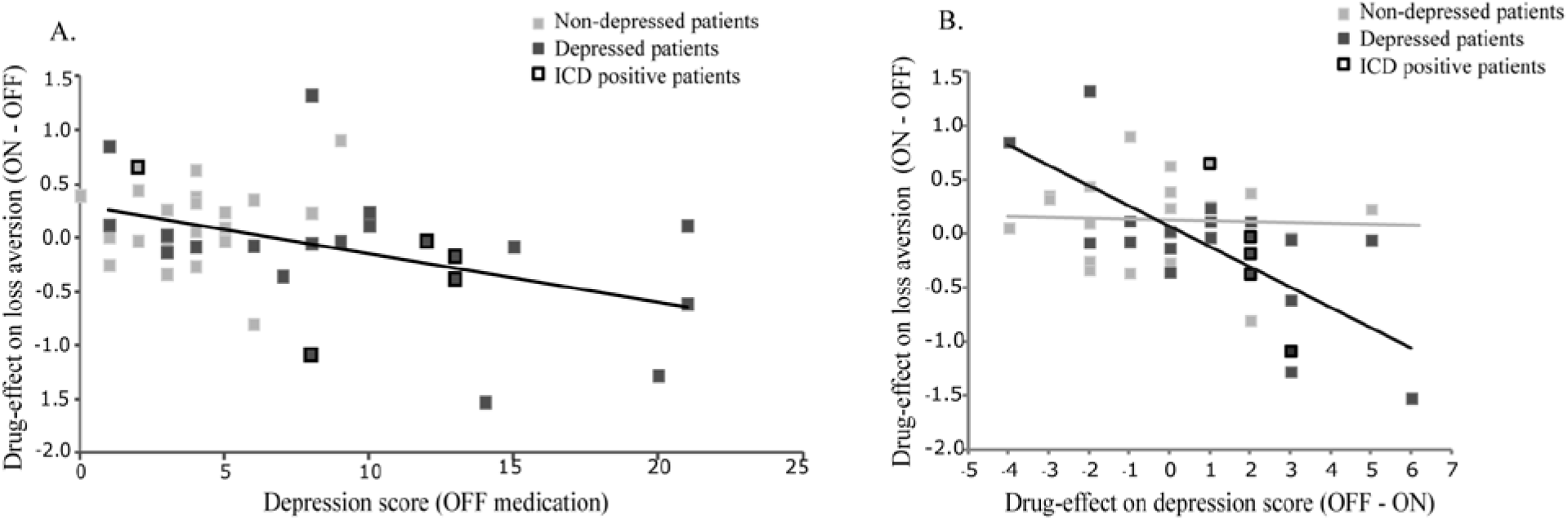
Correlations between (drug effects on) loss aversion and depression. **A** Correlation between scores on the Beck Depression Inventory during the OFF session (x-axis) and drug effects on loss aversion (λ) on the y-axis (ON session score minus OFF session score) (*rho*_(41)_=−.384, *p*=0.011). **B** Correlation between drug effects on depression scores on the x-axis (BDI score OFF session minus BDI score ON session) and drug effects on loss aversion (λ) on the y-axis (ON session score minus OFF session score). Depressed patients are marked in dark grey (*rho*_(19)_=−.592, *p*=0.005), non-depressed patients in light grey (*rho*_(20)_=−.021, *p*=0.93). This correlation was significantly different between groups (Fisher r-z transformation, *z*=−2.01, *p*=0.044). Patients who screened positive for having an ICD are marked with a black border.

In a supplementary analysis we compared PD patients, each group and drug session separately, with healthy controls. The median loss aversion parameter in healthy controls was significantly higher compared with non-depressed patients OFF medication (*U*=150, *p*=0.019), but not different from non-depressed patients ON medication and depressed patients during both the ON and OFF session (Table 2).

**Table 2.**
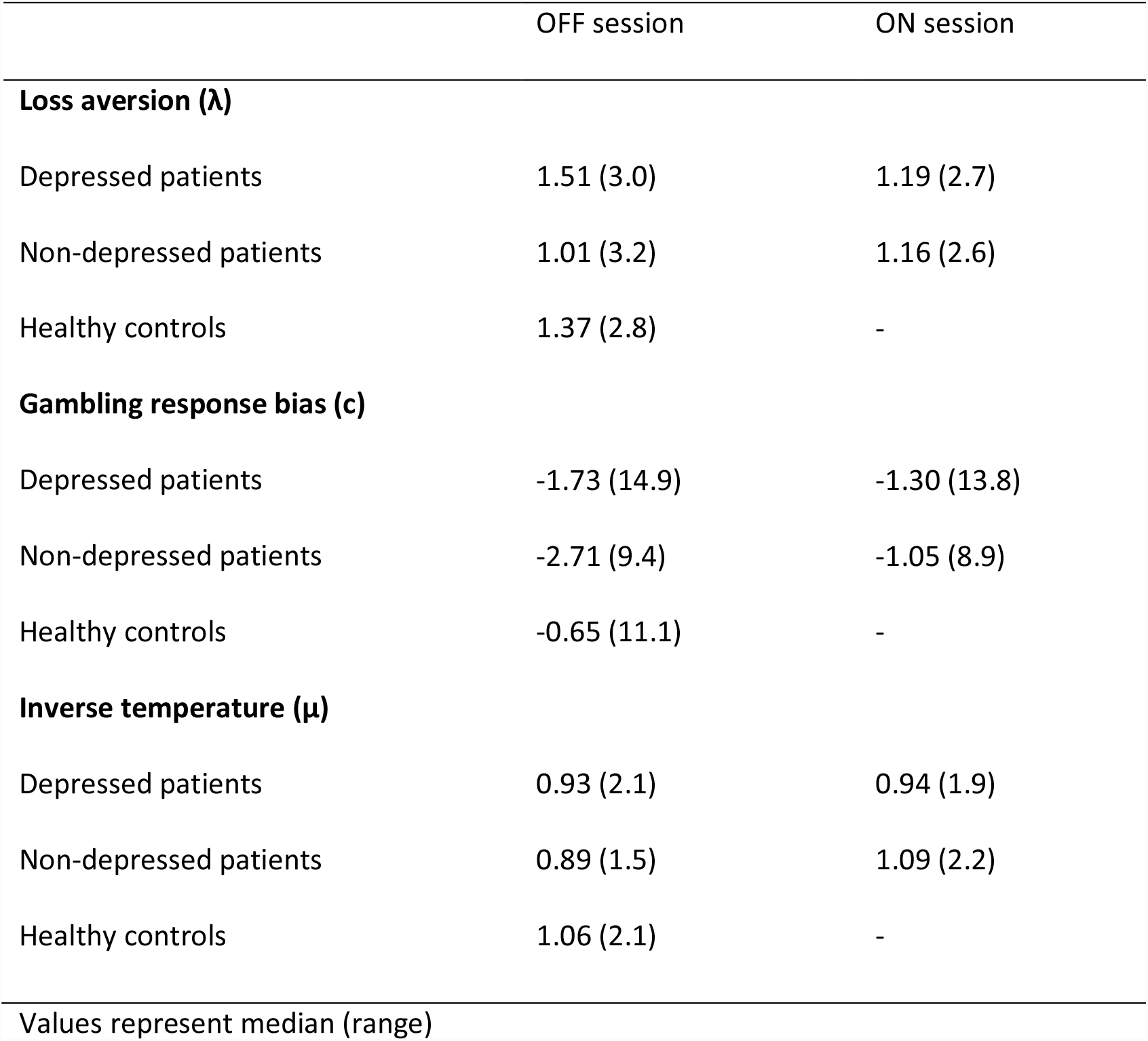
Model parameters per group and drug session

### Gambling response bias and inverse temperature parameter

The median gambling response bias and inverse temperature parameters are presented in Table 2 per group and drug session. These parameters were not normally distributed as indicated by Shapiro-Wilk test. We therefore used nonparametric statistics. Analyses of the gambling response bias parameter (c) revealed that there were no main effects of drug (*Z*=−1.78, *p*=0.08) or group (*U*=199, *p*=0.44), and no significant group*drug interaction (*U*=156, *p*=0.07). There were also no main effects of drug (*Z*=−.31, *p*=0.75) or group (*U*=225, *p*=0.88) and no significant group*drug interaction (*U*=226, *p*=0.90) on the inverse temperature parameter.

Relative to controls, non-depressed PD patients showed a significantly lower gambling response bias during the OFF session (*U*=111, *p*=0.001), but not during the ON session (*U*=177, *p*=0.08). By contrast, depressed PD patients showed a significantly lower gambling response bias during the ON session (*U*=151, *p*=0.033), but not during the OFF session (*U*=160, *p*=0.06) relative to controls. There were no differences in terms of the inverse temperature parameter (μ) between controls and either group of PD patients (ON and OFF medication).

### Proportion of accepted gambles

In addition to the computational parameters underlying risky choice, we analyzed the proportion of accepted gambles, which is a compound measure of risky choice. The proportion of accepted gambles in depressed PD patients was 53.6% OFF medication and 57.9% ON medication. In non-depressed PD patients this was 56.3% OFF medication and 60.2% ON medication. The proportion of accepted gambles in healthy controls was 62.2%. Mixed ANOVA in PD patients revealed no significant group*drug interaction (*F_(1,41)_*=0.01, *p*=0.94) and no main effect of group (*F_(1,41)_*=0.31, *p*=0.58) or drug (*F_(1,41)_*=2.32, *p*=0.136). The correlation between drug-induced increases in gamble acceptance and depression scores OFF medication failed to reach significance (*r*_(41)_=0.273, *p*=0.077). There was also no significant correlation between LED and drug-induced increases in gamble acceptance (*r*_(41)_=0.171, *p*=0.27). Comparison of patients with healthy controls (each patient group and drug session separately) revealed no significant differences in gamble acceptance.

## Discussion

The present study shows that dopaminergic medication induced differential effects on loss aversion during risky choice in PD patients with and without depression. Moreover, we demonstrate that the degree to which medication reduces loss aversion correlates with current depression severity and with drug effects on depression scores: drug-induced reductions in loss aversion were greater in more severely depressed patients and in patients who exhibit greater medication-related decreases in depression scores.

It is well known that dopaminergic treatment in PD patients can elicit detrimental side effects in the domain of risky choice. In experimental settings, dopaminergic medication increases risky choice in PD patients (Brand et al., 2004; Euteneuer et al., 2009), while also eliciting abnormal impulsive betting behavior during decision making (Cools et al., 2003). The present findings suggest that these effects might have been driven by patients with relatively higher depression scores in the OFF medication state. Critically, in those prior studies, the computational mechanisms underlying increased risky choice were not investigated. In this study, we adopted a computational approach, enabling us to isolate the mechanisms underlying drug-induced change during risky choice in (specific subgroups of) PD patients.

In line with results from a number of clinical trials in PD, we observed that dopaminergic medication significantly decreased depression scores in the depressed PD group (Barone et al., 2006; Barone et al., 2010; Stacy et al., 2010). Our data also revealed that dopamine-induced changes in depression scores correlated with dopamine-induced changes in loss aversion. Patients with the greatest antidepressant effect of dopaminergic medication also exhibited the greatest decrease in loss aversion. These findings raise the hypothesis that dopamine-induced changes in loss aversion might underlie the beneficial effects of dopaminergic medication on depressive symptoms in PD.

In prior work, we have put forward the dopamine overdose hypothesis to account for the detrimental effects of dopaminergic medication on punishment-based learning and decision-making. This hypothesis states that dopaminergic medication doses necessary to remedy dopamine levels in severely depleted dorsal striatum might detrimentally overdose dopamine levels in the relatively intact ventral striatum (Cools et al., 2001, 2003; Cools, 2006). According to this hypothesis, one might expect that any abnormal decrease in loss aversion is seen only in non-depressed PD patients with a putatively intact ventral striatum, while not extending to depressed PD patients, who have been argued to exhibit ventral striatal dopamine deficiency (Vriend et al., 2014). By contrast, the current study suggests that depressed PD patients are particularly at risk for developing detrimental effects of dopaminergic medication on risky choice. This finding concurs generally with clinical evidence indicating that PD patients who exhibit more severe depressive symptoms are at increased risk for having ICD (Pontone et al., 2006; Isaias et al., 2008; Voon et al., 2011b), although a strong link between (dopamine-induced decreases in) loss aversion and ICD has yet to be established (Voon et al., 2011a; Giorgetta et al., 2014).

Neuroimaging studies with healthy and depressed individuals (without PD) have revealed neural loss aversion in several limbic brain regions, including the striatum (Tom et al., 2007; Canessa et al., 2013; Chandrasekhar Pammi et al., 2015). Moreover, evidence from work in healthy volunteers, PD patients and rodents indicates that both drug-induced impulsivity and risky choice as well as depression are accompanied by low striatal dopamine D2 receptor and dopamine transporter (DAT) availability (Remy et al., 2005; Weintraub et al., 2005; Boileau et al., 2009; Buckholtz et al., 2010; Cocker et al., 2012; Norbury et al., 2013; Vriend et al., 2013). Together these observations raise the hypothesis that drug-induced changes in loss aversion in PD could reflect impaired auto-regulation of striatal dopamine levels. This hypothesis might be tested in future studies by combining the use of neuroeconomic tools and controlled medication withdrawal in PD, with neurochemical imaging of dopamine D2 receptor availability and dopamine release.

A number of limitations of the current study should be highlighted. First, in contrast to recent findings from (Chandrasekhar Pammi et al., 2015), who observed greater loss aversion in depressed patients (without PD), we did not observe significantly greater loss aversion in our depressed PD patients OFF (or ON) medication than controls. This might reflect the fact that not all our patients were currently depressed. Instead we included patients based on the presence of a history of depression. This should be tested in future work with a larger sample of currently depressed patients. Second, the design was not optimized for comparing patients with controls, who were tested only once OFF medication, in contrast to the patients, who were tested twice, once ON and once OFF medication. Moreover, our control group exhibited a median loss aversion parameter of 1.4, which is relatively low compared with previous studies with healthy individuals in which loss aversion estimates ranged between 1.4 and 2.0 (Tom et al., 2007; Sokol-Hessner et al., 2009; De Martino et al., 2010). This apparent discrepancy with prior work might reflect the fact that our control group was somewhat older, but in any case caution is warranted when interpreting the results from the comparisons with healthy controls. Finally, the pattern of medication effects in terms of the proportion of accepted gambles, which did not exhibit a significant drug*group effect, is quite different from that in terms of the loss aversion parameter, which did exhibit a significant drug*group effect. This is not surprising because our model takes into account the prior theoretical insight that the proportion of accepted gambles is a function of multiple parameters, including not just loss aversion, but also gambling response bias. In fact, effects of medication and depression on loss aversion were isolated, precisely because we disentangled it from any (non significant) variability in terms of gambling response bias. As such, the discrepancy between pattern of effects on the proportion of accepted gambles and that on loss aversion highlights the strength of the adopted modeling approach.

## Acknowledgements

This project was funded by a grant from the “Stichting Parkinson Fonds”, Hoofddorp, the Netherlands. We would like to thank all participants for their cooperation in the study.

